# Dynamics and maintenance of categorical responses in primary auditory cortex during task engagement

**DOI:** 10.1101/2022.12.19.521141

**Authors:** Rupesh Kumar Chillale, Shihab Shamma, Srdjan Ostojic, Yves Boubenec

## Abstract

Grouping sets of sounds into relevant categories is an important cognitive ability that facilitates the association of stimuli with appropriate goal-directed behavioral responses. In perceptual tasks, a prominent role of primary auditory cortex (A1) is to multiplex the encoding of sound sensory features and task variables. Here we investigated the involvement of A1 in initiating sound categorization. We trained ferrets to discriminate click trains of different rates in a Go/No-Go delayed categorization task and recorded neural activity during both active behavior and passive exposure to the same sounds. Purely categorical response components were extracted and analyzed separately from sensory responses to reveal their contributions to the overall population response throughout the trials. We found that population-level representation of Go/No-Go behavioral categories emerged during sound presentation and was present in both active behavioral and passive states. However, upon task engagement, categorical responses to the No-Go category became suppressed, leading to an asymmetrical representation of the Go stimuli relative to the No-Go sounds and prestimulus baseline. At stimulus offset, the population code changed abruptly but categorical representation was maintained during the delay period. Categorical responses extracted from the stimulus period correlated with the ones found in the delay epoch, suggesting an early contribution of A1 to stimulus categorization.

## Introduction

Grouping individual stimuli into abstract, context-dependent categories and maintaining them in memory are fundamental cognitive abilities for appropriate behavioral responses and decisions. Neural correlates of such categorical perception have been found widely across cortical areas and modalities, from sensory cortices to the frontal regions^1–4^. The emerging picture thus far has been that categorization is a hierarchical process implemented through a series of computations and gradual transformations across multiple cortical areas, beginning with relatively basic stimulus representations in the primary sensory areas, and ultimately concluding with categorical responses in the frontal regions^3,5,6^. In the auditory modality, this framework suggests that the primary auditory cortex (A1) would mainly represent stimulus acoustic features, and only exhibit weak correlates of the associated behavioral categories^7–9^.

Recent studies however have challenged this conception from multiple viewpoints. To begin with, the encoding of sounds in the primary auditory cortex (A1) is relatively plastic, and is rapidly enhanced by task engagement^3,10–12^. Furthermore, A1 has already been shown to extract some of the behavioral meaning of target sounds for task-relevant downstream readout^13–15^. Nevertheless, the stimuli and experimental designs used on most of these experiments didn’t allow for disentangling category-specific neural dimensions from sensory-related activity. It thus remains uncertain how A1 population responses represent stimulus category during sound presentation in addition to strictly sensory-evoked responses, and how the representations of the task-relevant categories could be dynamically maintained after the stimulus is played. This study explores the critical questions of how and when categorical encoding emerges in primary auditory cortex, and the extent to which these categorical representations become behaviorally-shaped upon task engagement.

To this end, we trained ferrets to classify click trains into Go and No-Go categories of either low or high rates during an appetitive Go/No-Go task. After training, we recorded in A1 while ferrets passively listened to or actively discriminated between the two stimulus categories. Using population-level analyses, we contrasted the representation of stimulus features (click-rates) and behavioral categories (Go/No-Go) during and after stimulus presentation. First, we discovered a categorical encoding emerging early during stimulus presentation. Second, upon task engagement, the representations of the No-Go category became suppressed, leading to an asymmetrical representation of the Go stimuli relative to the No-Go sounds and prestimulus baseline. Third, at stimulus offset, the population code changed abruptly but the amplitude of the categorical response persisted beyond the stimulus period to the delay epoch. The categorical responses built up during the delay in anticipation of the subsequent behavioral response. Last, incorrect behavioral choices could be traced back to degraded sensory encoding during the stimulus period, resulting in a degraded categorical representation.

## Results

### A1 neurons sustain activity after Go sounds during task engagement

Two ferrets were trained on a Go/No-Go delayed categorization task under appetitive reinforcement. Water-deprived ferrets had to classify click trains into two categories: target (Go) and non-target (No-Go) depending on the rates of click trains. Six rates were used, from 4 to 24 Hz in 4 Hz steps, and with a category boundary fixed at 14 Hz. To ensure the dissociation between categories and stimulus rates, one animal was trained with low rates as the Go sounds, while the second animal classified high rates as the Go sounds. Click trains were presented for 1.1 s, and were followed by a 1 s-long delay in which the animals had to refrain from licking (Fig 1a). Licks during the subsequent 1 s-long response window were rewarded with water in Go trials, and punished with a timeout in No-Go trials. Any licks before this response window (Early licks) resulted in an aborted trial and were punished with a timeout; these were called “Early trials”.

**Fig 1.**
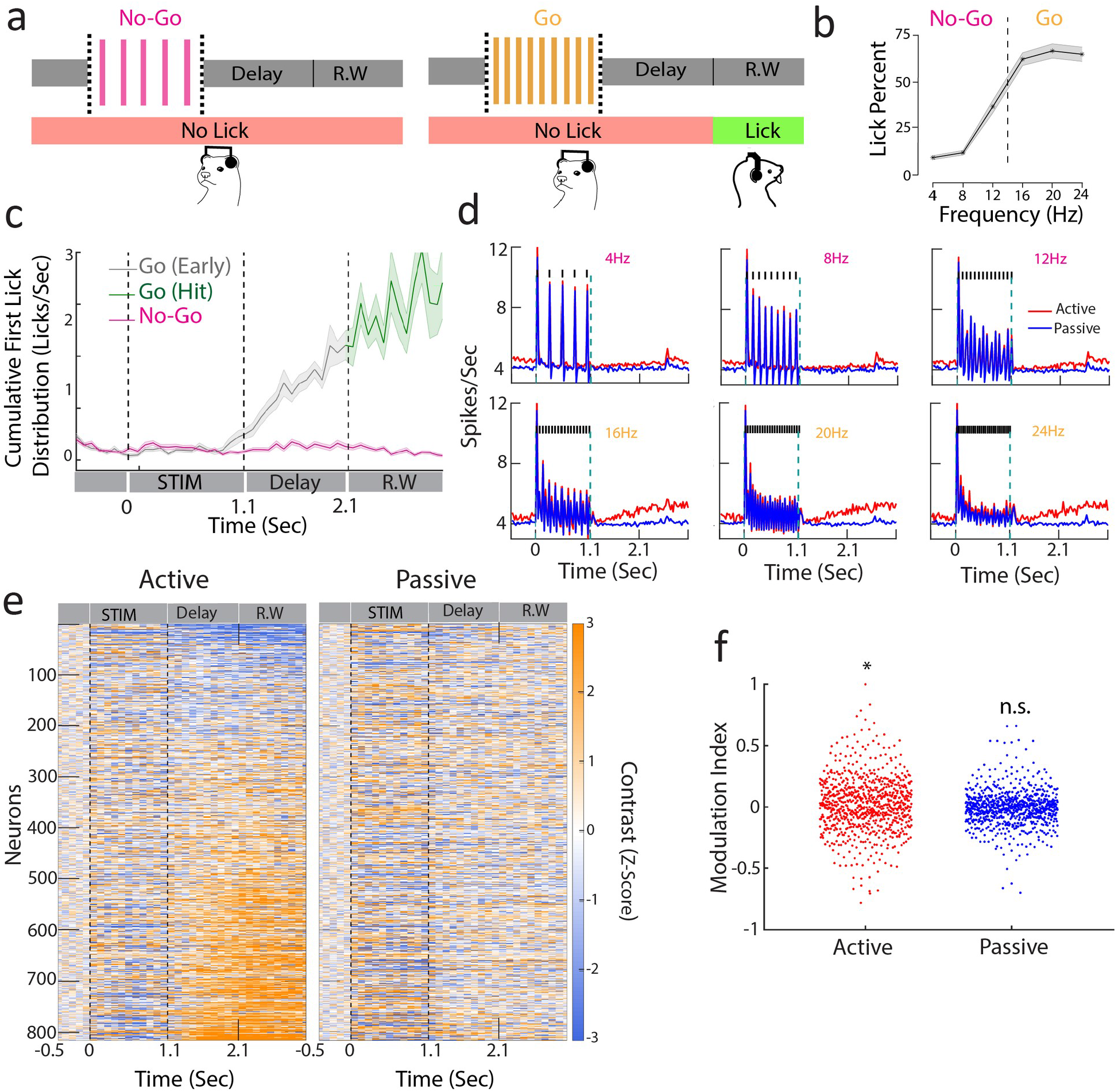
Sustained A1 activity during a delayed categorization task. **a.** Delayed categorization task. A trial starts with a 0.5 sec of pre-stimulus silence followed by 1.1s duration click train stimulus. The animal must wait for a 1s delay period before the 1s long response window. Correct trials were rewarded with water while error trials and early trials (lick during delay period including sound period) were punished by a timeout. **b.** Proportion of licks in response window for one of the animals (Ferret P) with low rates as No-Go and higher rates as Go stimuli. Only non-early trials are considered. **c.** Temporal profile of first lick rate. **d.** Average PSTHs of all neurons corresponding to passive (blue curve) and active (red curve) states for each of the click trains (only correct behavioral trials were used, i.e. correct rejections for No-Go and hits for Go sounds). Note that the response during the delay period is enhanced for Go stimuli (16, 20, 24 Hz). **e.** Contrast (Go – No-Go firing rate) computed for neurons of both animals (n=816 units) in passive (right) and active (left) states (z-scored with pre-stimulus baseline activity). Neurons were ranked by delay firing rate for Go active trials. **f.** Modulation Index (see Methods) unit for delay activity for active and passive states (t-test * p<0.05; n=816 units).

Ferrets categorized the click trains (Ferret P: d’ = 1.2±0.5 n=35 sessions; Ferret T: d’ = 1.1±0.2 n=39 sessions; Fig 1b) with a bias to lick for the No-Go rates close to the category boundary (12 Hz for the animal shown in Fig 1b). This was found in both ferrets (Fig S1a,b) regardless of the mapping between stimulus rates and category, suggesting that their decision criterion was relatively liberal and impulsive. First lick probability was higher in Go than in No-Go trials throughout the entire trial-duration (Fig 1c), confirming the categorical response profile. We also observed a build-up of lick probability during the delay, with Early responses being quite common in Go trials. Early trials and error trials (misses and false alarms) were discarded in the analysis unless otherwise specified.

We chronically recorded neural activity from a total of 816 units in A1 (Ferret P: 575, Ferret T: 241), while the two animals alternately listened passively to the stimuli or actively engaged in categorizing them. Figure 1d depicts the average click responses across all cells, showing a significant increase in firing rate relative to baseline activity in all conditions. Firing rates also steadily ramped up during the delay period in Go trials (Fig 1d,e; both animals in Fig S1c,d). Importantly here we considered only Hit trials in which the animals were not licking the spout until the response window. Neurons with significant changes in firing rate during the delay period did not show any modulation of activity during the passive state or after the No-Go sounds, reflecting the maintained representation of the Go sound category in the task-engaged delay period. Additionally, we found that these neurons did not uniformly increase their firing rates during the delay period, but instead exhibited heterogeneous trends with a bias towards excitatory responses during the delay (Fig 1f t-test active p=0.01, passive p=0.15, n=816 units; both animals in Fig S2 Ferret P: active p=0.04, passive p=0.83, n=575 units; Ferret T: active p<0.001, passive p=0.35, n=241 units).

### A1 population activity encodes behavioral categories during stimulus in both active and passive states

Because A1 activity was heterogeneously modulated during the post-Go sound delay, we relied on population decoding to examine the categorical representation. To do so, we trained linear decoders to discriminate Go and No-Go trials based on single-trial population activity (Fig 2a). During task engagement, population responses steadily discriminated between Go and No-Go categories along the entire course of the trial (Fig 2a, both animals in Fig S3). Decoding accuracy diminished at stimulus offset, but subsequently increased during the delay period, ultimately peaking in the behavioral response window (Fig 2a and Fig S3). By contrast, accuracy in the passive context decreased throughout the delay period and was similar to what is observed in the naive animal (Fig S4).

**Fig 2.**
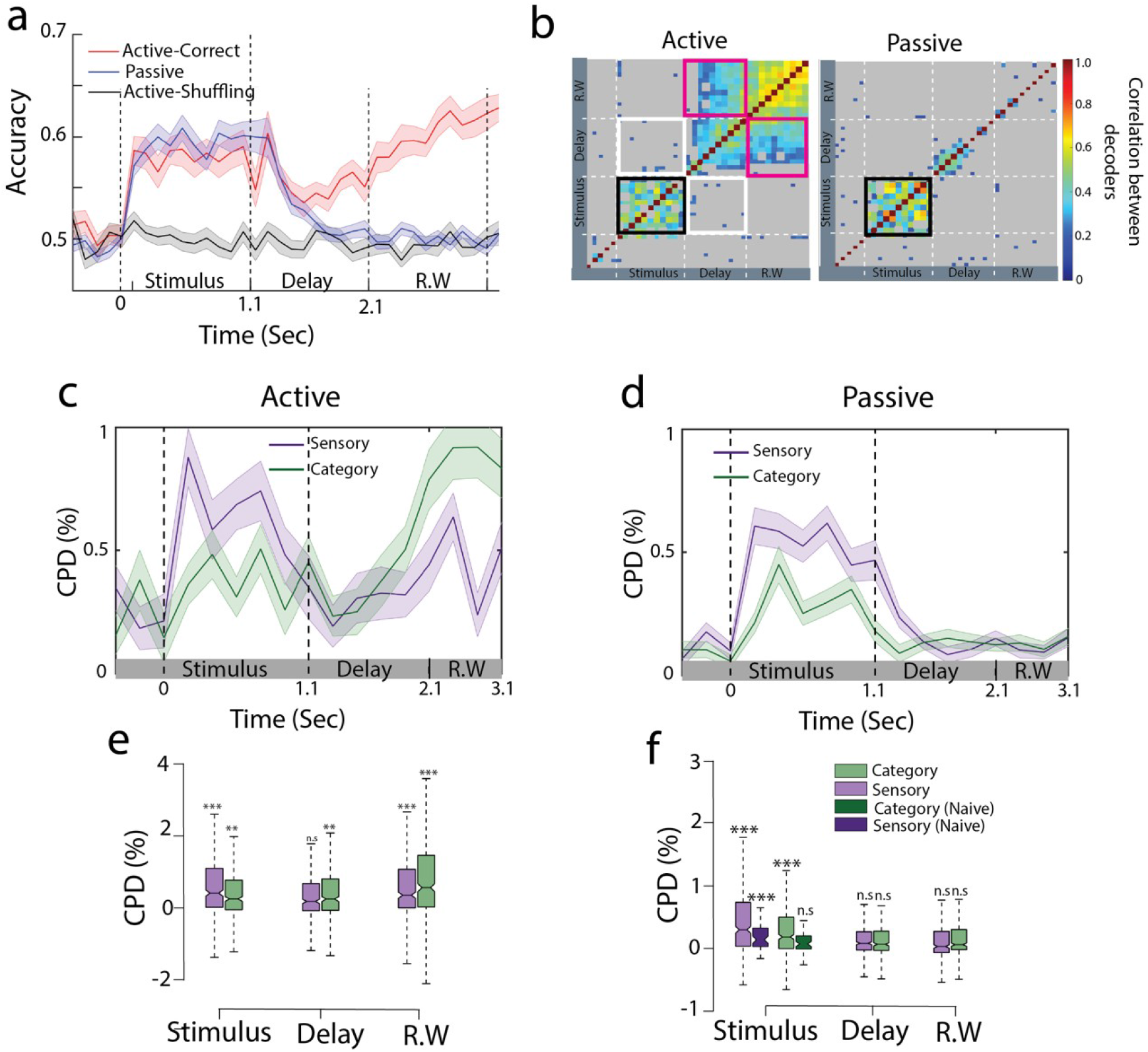
Population decoding of categorical information from primary auditory cortex. **a.** Go vs No-Go classification performance in active (red curve) and passive (blue curve) conditions (n=35 sessions). Gray curve indicated the performance by shuffling labels for the task-engaged condition. Error bars show 1STD. **b.** Cross-correlation matrix between decoders trained at different time points for passive (left) and task-engaged (right) data. Non-significant correlations are shown in grey. Significance was assessed by permutation test (200 permutations). Black frame shows the significant cross-correlation during the stimulus period. White (resp. pink) frames show absence of correlation between decoders trained during the stimulus and the delay (resp. response window) period. **c.** Coefficient of partial determination (CPD) computed by fitting linear regression models in active state. Shaded region represents 1 SEM over all the neurons. **d.** Same as **c** for passive state. **e.** CPD computed during the stimulus, delay and response window time epochs. Significance is tested against pre-stimulus period value (two-tailed t-test *** p<0.001, n=395 neurons). **f.** Same as **e** for passive state. CPD computed from a naive animal is added in the stimulus period for comparison.

We then looked further into how population code evolves during the trial. For this, we compared decoders trained at different time bins throughout the trial. We found that for both passive and task-engaged sessions, the direction of the decoding axis was mainly preserved during the stimulus epoch (black outline in Fig 2b), and the corresponding decoders trained on passive and task-engaged data were correlated (Fig S5). Furthermore, these decoders were uncorrelated with the decoding axis during the *delay* period of the task-engaged condition (white outline in Fig 2b), indicating a behavior-dependent change of the population code at the sound offset. This delay activity pattern further persisted during the *response window*, as shown by the correlation between delay and response window decoding axes (pink outline in Fig 2b). This suggests a time-independent categorical representation in A1 population activity throughout the delay and response window epochs.

While categorical encoding is evident during task engagement *after* the stimulus end, it remains unclear whether it was biased towards stimulus-related or category-related responses *during* the stimulus. Indeed, both stimulus-related (click train rate) and category-related responses are captured by the decoders (Fig S6a,b). To disentangle sensory and categorical contributions, we computed a population-level linear regression of the neuronal firing rates at each time bin in each session (see Methods for details). Two regressors were used: (i) stimulus rates as a *sensory* regressor, and (ii) the binary category (Go or No-Go) as a *category* regressor. We also fitted two additional models in which the labels of one of the regressors were shuffled^16^, which allowed us to compute the relative contributions using a *coefficient of partial determination* (CPD) from each regressor as the decrement in variance due to shuffling one of the regressors. During task engagement, we found that the CPD of the category regressor increased during the stimulus period and persisted throughout the delay period, before increasing at the response window (Fig 2c,e; paired t-test on category regressor, delay vs. response window; Ferret P n=395 neurons p<0.001; Ferret T n=203 neurons p<0.01). Interestingly, both task-engaged and passive category CPD were larger than chance during stimulus presentations (Fig 2c-f), an effect absent in the naive animal (Fig 2f), which indicates training-dependent encoding of categories.

### Population-level categorical responses shows suppression of No-Go representations upon task engagement

We then examined how the category information during the delay emerged from a mixed representation of stimulus and category during stimulus presentation. Regression axes define encoding axes along which population activity carries information about stimulus click rate or category. We thus projected population activity onto the category regressor axis to track the categorical representation in the population at each moment in time. This procedure sums the neuronal responses weighted by the coefficients extracted from the earlier regression analysis and reduces A1 population dynamics to one information-bearing dimension encoding category through time, independently from sensory information. We did so in both the stimulus and the delay periods for the passive and task-engaged conditions in order to compare the patterns of categorical encoding across conditions.

Projections on the category regressor (called thereafter categorical responses) revealed a stable encoding of categories during sound presentation, regardless of the stimulus identity (Fig 3a,b), indicating that these categorical responses were not contaminated by sensory activity. We measured the magnitude of the categorical responses with a category index measuring the distance between the projections of the rates at the boundary (12 and 16 Hz) minus the average distance between all the pairs of adjacent rates within categories (Fig S7a,b). We found that categorical indices were identical between passive and task-engaged states during stimulus presentation (Ferret P: p=0.80 n=35 sessions; Ferret T: p=0.78, n=39 sessions paired sample t-test), in contrast to the large difference found during the delay period (Fig 3c-e).

**Fig 3.**
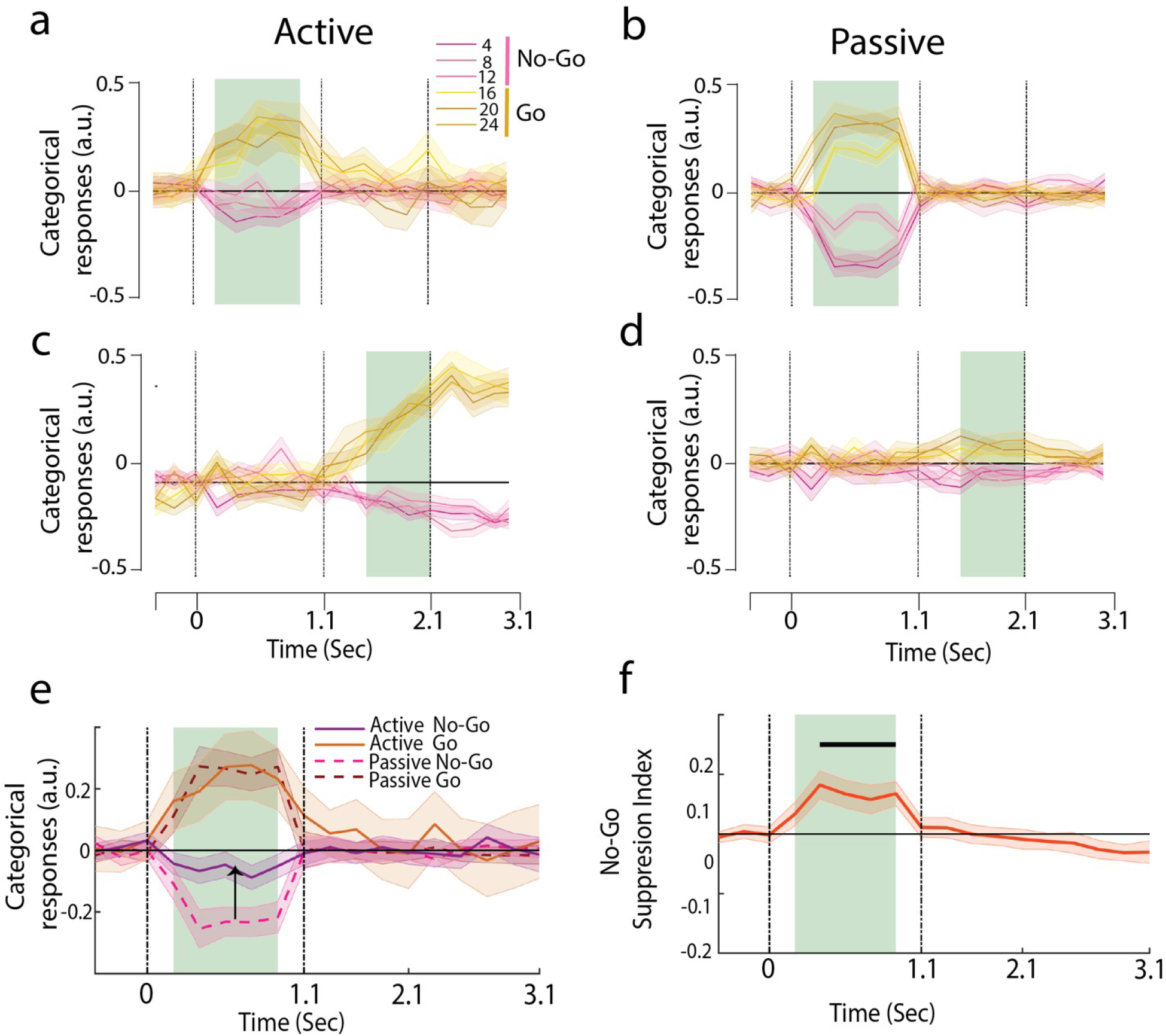
Emergence of categorical representation through suppression of No-Go sounds. **a,b,c,d.** Projection of trial averaged activities of individual click rates onto category regressors trained at different time epochs. The shaded regions show training time and the error bars are ± 2 SEM over sessions. **a,c** are for active and **b,d** are passive states. See Supplementary figure S8 for graded sensory responses to the different stimuli. **e.** Categorical responses for passive and active states. **f.** Time-course of No-Go suppression in categorical responses.

To further determine if task engagement induced a targeted change along the categorical neural dimension, we examined the temporal profiles of the Go and No-Go categorical responses. Here we found that there was a clear shift of population coding between the two behavioral conditions: No-Go categorical responses aligned with the pre-stimulus spontaneous activity during task engagement (Fig 3a), inducing an asymmetry between No-Go and Go categorical responses (Fig 3e, both animals in Fig S7c,d), a characteristic that is not observed in the passive projections (Fig 3b). A No-Go suppression index measuring the relative displacement of the No-Go responses closer to the projections of spontaneous activity (Fig 3f) showed that the suppression was confined to the stimulus period (Fig S7e,f bottom row; Ferret P p<10^-6^ n=35 sessions; Ferret T p<10^-4^ n=39 sessions), and absent in the delay period (Ferret P p=0.70; Ferret T p=0.46). Crucially, projections of the passive population responses onto the active category regressor *did not* show the suppression of No-Go sounds, confirming that this shift originated from a change in the structure of the population responses during task engagement, and not due to the weights of the regressor themselves (Fig S9). In summary, these findings demonstrate a task-induced asymmetry in categorical representations which specifically targets the relevant neural dimension.

### Single-trial categorical representations correlate between sound and delay periods

The categorical signal in the stimulus period exhibits short latencies (Fig 3a) consistent with a feedforward mechanism of early categorization taking place in A1. We wondered whether this early categorical information may influence later categorical information captured in the delay period. However, the categorical axes in the early stimulus and delay periods were not correlated (white frames in Fig 4a). So how could the categorical information present in these two distinct intervals be still linked? To find out, we tested *directly* whether the categorical responses during the stimulus period correlated with those during the delay. Specifically, we examined carefully the *single-trial* fluctuations of categorical responses in each period and discovered that the two categorical responses were correlated (one session in Fig 4b, all sessions in the inset; p<0.001 permutation test n=74 sessions). Note that the stimulus and delay categorical responses were computed from uncorrelated projection axes (Fig 4a) that were originally derived from distinct non-overlapping epochs, and hence the single-trial fluctuations in categorical responses need not have been correlated in any way.

**Fig 4.**
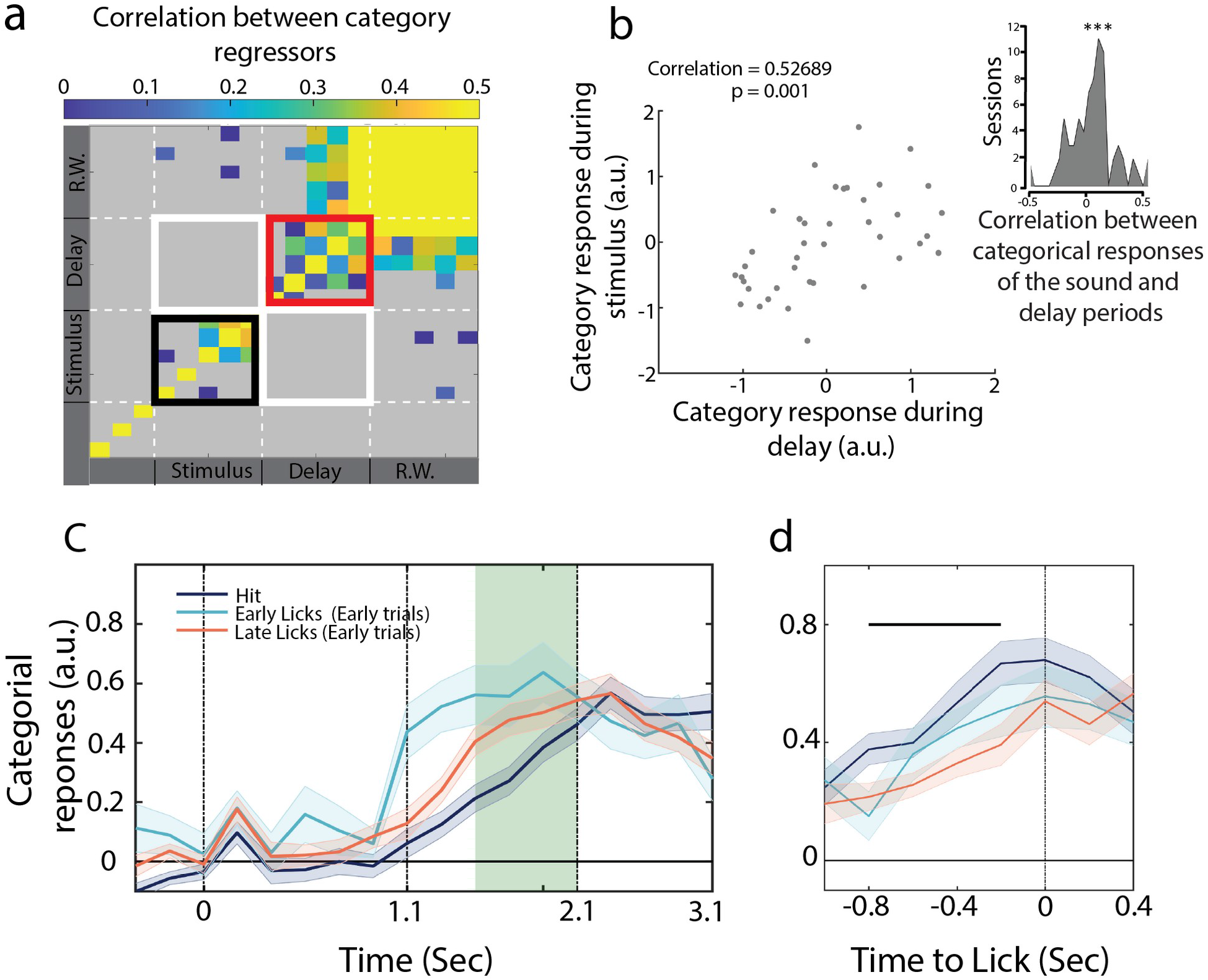
Post-stimulus anticipatory activity correlates with categorical representation during the sound. **a.** Temporal evolution of the category regressors. **b.** Scatter plot for categorical response during stimulus and delay period for one session. Inset: Correlation of single-trial categorical responses between the sound and delay periods (n=74 sessions). **c.** Categorical responses for Hit, Early trials with licks in the early (Early licks) and late (Late licks) phases of the delay period. **d.** Lick-aligned categorical responses. Projections were not different between Hit and Early trials at lick time (see trial decoding in Fig S11).

### Post-sound anticipatory activity builds up towards behavioral response

Another interesting relationship is that between the categorical responses and the licks. We have observed that population activity on hit trials ramped up throughout the delay period (Fig 3c and Fig 4c, categorical responses during the delay). By aligning singletrial projections onto lick timings during the response window, we found that the categorical activity built up during the delay period and culminated when the animal licked (Fig 4d, categorical responses centered on licks), indicating that delay activity was anticipating behavioral responses. We further tested whether the temporal dynamics of the categorical responses correlated with the timing of the animal’s behavioral response. To do so, we took advantage of early Go trials when the animals licked *before* the response window (see lick histogram in Fig 1c), so that we accessed to Go trials in which first licks occurred earlier than what we had in hit trials. We then sorted all the collected trials into three groups: early trials with licks early in the delay period (first half), early trials with licks late in the delay (second half), and hits. We found significantly faster response build-up rates in early trials than in hit trials (Fig 4c). Early trials with early licks in the delay exhibited the steepest response buildups (Fig 4c), a pattern that was not observed in the categorical responses during the stimulus period (Fig S10). We then aligned all trials to their respective reaction times and found that categorical responses of early and hit trials culminated at the licking time (Fig 4d). This alignment was done on lick times, i.e. independently of neural activity, indicating a buildup of population-level dynamics anticipating behavioral response.

### Sensory information is degraded during error trials leading to poor categorization

Finally, we searched for the neural correlates of incorrect perceptual decisions. In particular, we wanted to pinpoint which encoding stages were degraded during incorrect categorizations. We envisioned three different hypotheses causing the error trials: (a) sensory information was intact in error trials, but conversion to the correct category during the stimulus was incorrect; or that (b) sensory-to-category processing during the stimulus period is correct, but the anticipatory activity is impaired during the delay; or that (c) sensory information was in fact degraded early on during the stimulus, leading to a loss of categorical information.

We first sought to test whether the error trials correlated with an inversion of the neural responses between the Go and No-Go categories, as would be predicted by hypothesis (a). To do so, we used Go/No-Go classifiers trained with correct trials (as in Fig 2a) and reported the classifiers’ performance in predicting the expected (and not actual) behavioral categories. If false alarms were fully behaving like hits, and misses like correct rejections, we should observe below-chance decoding accuracy. That was not the case, but instead we found a decrease in decoding accuracy during the sound period, and a decoding performance at chance level during the delay (Fig 5a,b; decoding accuracy during delay for incorrect trials: p=0.99, permutation test). As a control, we noted that decoding accuracy dipped below chance during the response window confirming that motor information was confined to this late time period (Fig 5a). This indicates that incorrect decisions could be traced back to the stimulus period, excluding hypothesis (b) in which only the delay activity was incorrect during error trials.

**Fig 5.**
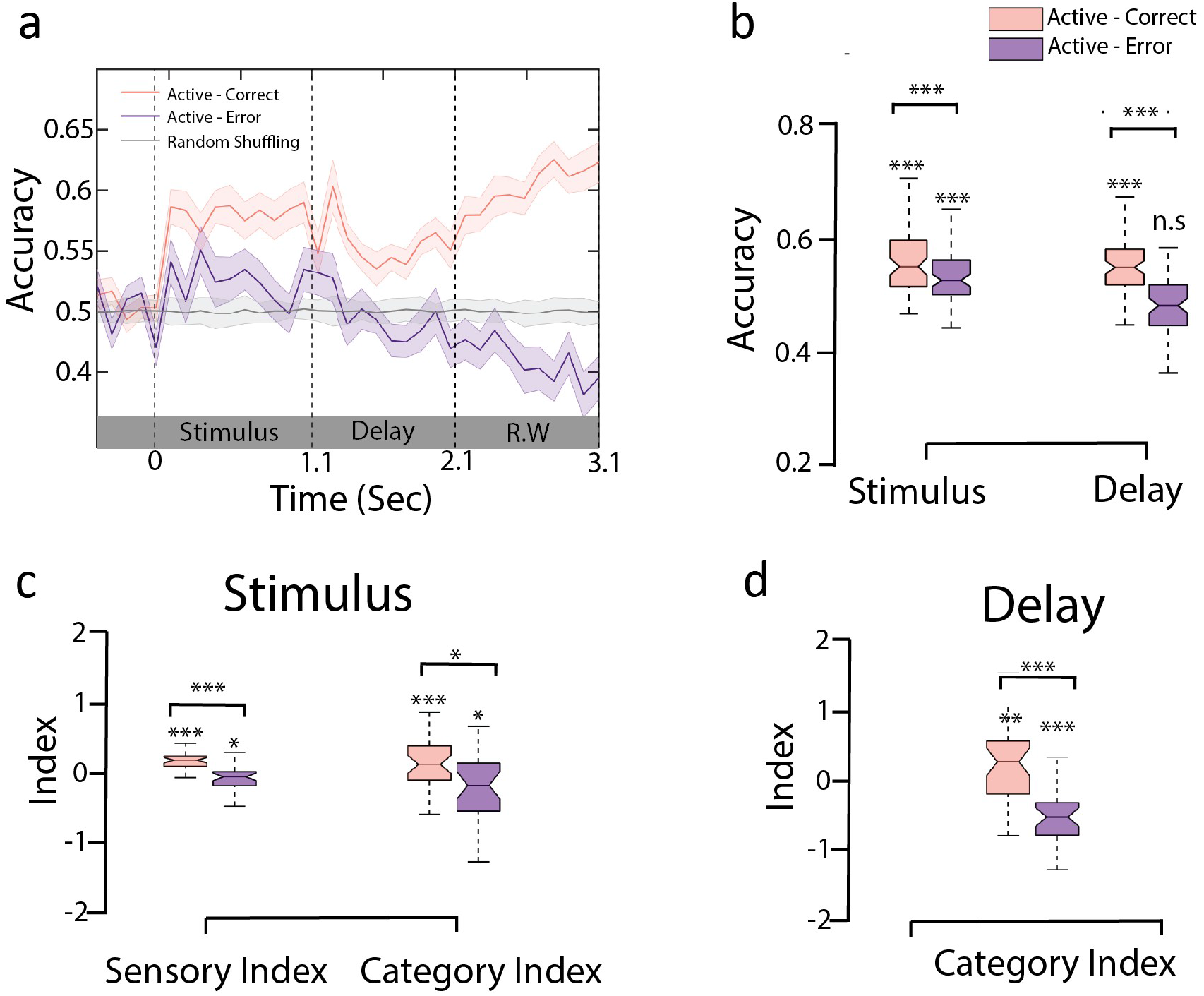
Sensory and categorical responses during error trials. **a.** Encoding of behavioral choices: here we trained the active classifier on correct behavioral choices (Hit and Correct Rejection) and used the decoding weights to compute the accuracy using incorrect behavioral choices (False alarm and Miss) as shown in the cyan curve. Gray curve indicated the performance by shuffling labels in task engagement. Error bars show 1STD. **b.** Accuracy of the active decoder for correct and error trials during sound and delay period **c.** Categorization index and sensory index computed from the projections of trial averaged correct and error trials onto category regressor trained during sound period. **d.** Same as **e**, for delay period.

We then further tested the hypothesis (c) of a degradation of sensory information during the stimulus period. In line with this interpretation, we found that both sensory and categorical responses were degraded in incorrect compared to correct trials (Fig 5c and Fig S12a,b for both animals; stimulus period-SI: Ferret P p<10^-5^ n=35 sessions; Ferret T p<10^-3^ n=39 sessions; stimulus period-CI: Ferret P p=0.01 n=35 sessions; Ferret T p=0.35 n=39 sessions; paired t-test across sessions, individual boxes were tested with permutation test). Categorical responses were further degraded during the delay period (delay-CI Fig 5d and Fig S12b,c for both animals; paired t-test across sessions; Ferret P p<10^-4^ n=35 sessions; Ferret T p<10^-5^ n=39 sessions). Altogether this suggests that errors likely originated from an improper encoding of the stimulus that subsequently led to an incorrect categorization and behavioral response.

## Discussion

### Early build-up of categorical encoding in A1

Categorical perception of real-world signals is a key cognitive function in all sensory modalities, one that is thought to be implemented at higher cortical levels^1,3,17–19^. In this work we examined if and how population responses in *primary* auditory cortex could contribute to and reflect the categorical encoding of sound during passive listening or engagement in a Go/No-Go categorization task. The study resulted in four main findings. *First*, we isolated an encoding of behavioral categories during stimulus presentation in both task-engaged and passive listening. This was not observed in a naive animal. We interpret this observation as an effect of long-term memory^20^. *Second*, we found that categorical responses to No-Go sounds were suppressed during task engagement during the stimulus period. *Third*, the categorical representation changed at stimulus offset *but still* correlated at the single-trial level with categorical responses observed during the stimulus period. *Fourth*, incorrect behavioral choices were traced back to degraded categorical encoding during the *stimulus period*, resulting in a degraded categorical representation during the subsequent *delay* period. Altogether these results suggest that behavior-dependent categorical information emerged during the stimulus period, influenced population dynamics beyond the stimulus period itself, and persisted throughout the delay period until the behavioral response.

Previous findings have found categorical responses in A1 with a marked increase of response contrast at category boundaries^3,12^. Here we focused on revealing the temporal dynamics of categorical encoding at the population level, and how it changed after stimulus offset. Regression analyses allowed us to disentangle the binary categorical representation from the more graded representation of the sensory properties (i.e., click train rates) (Supplementary figure S8). Sensory responses exhibited shorter latencies than categorical ones, in line with the idea that the internal dynamics in A1 contribute to the feedforward conversion of sensory information into behavioral categories.

### Population-level suppression of No-Go categorical responses

Our analyses showed that, even though categorical encoding was present in passive and active states in trained animals, task engagement induced a suppression of No-Go responses during the stimulus. This was not due to a lack of responses to No-Go sounds, as shown by the average population PSTH (Fig 1d) and its sensory responses (Supplementary Fig 13). Instead, this asymmetry in the categorical representation stems from a task-induced population-level enhancement of Go and suppression of No-Go categorical responses, consistent with our previous findings^14^ in these Go/No-Go paradigms. To elaborate, No-Go sounds instruct the animal to maintain the same behavioral output (lick inhibition) as in periods of silence when spontaneous activity is measured. We therefore proposed that the alignment of the No-Go population responses with the spontaneous activity reflects the identical behavioral meaning of these two epochs. Here we found a similar mechanism (suppression of No-Go responses) which was confined to the categorical responses extracted through linear regression. This suggests an interesting mechanism in which population dynamics could multiplex behavior-independent sensory representations and task-modulated categorical encoding. Intriguingly, this is reminiscent of population dynamics found in prefrontal cortex (PFC)^21^, where sensory information and decision formation were represented along different neural dimensions.

In line with a role of A1 in the category build up, we found that categorical information was degraded in error trials, which is consistent with an early mapping of individual stimulus into generic categories at the level of primary sensory areas. We thus propose that A1 has an initial contribution in the feedfoward formation of behavior-dependent categories. During behavior, higher areas would access an explicit representation of the behaviorally-relevant categories by reading out the asymmetrical population-level encoding of Go and No-Go sounds in A1^14^. These areas would then utilize the A1 categorical representation to amplify and strengthen the ongoing category, possibly passing or gating it to motor-related regions depending on the behavioral state^5^.

### Possible roles for post-sound sustained activity

Categorical responses changed at stimulus offset to maintain a prolonged activity during the delay period. We do not interpret this activity as efferent copies directly sent from motor regions^22^, as one would expect such motor-related activity to have a short-latency (~100 ms^14^). Here, sound category was successfully decodable throughout the entire delay period (Fig 2a), which is counter to the functional implications of short-latency efference copies. We have also found that the delay activity is not tied to licks since false alarm trials, in which the animals licked during the response window, were not decoded as hit trials (Fig 5b). Therefore, one possible conclusion is that the delay activity in A1 is the result of a feedback signal from higher-order decision-related areas, engaging A1 in a network of parietal and frontal areas^23^. Consistent with this hypothesis, we have demonstrated that trial-to-trial fluctuations of categorical responses during the *sound* period correlate with the amplitude of the *delay* categorical signal (Fig 4b). This is reminiscent of rotation in population representations recently found in V1^24^, where rotational dynamics played a role in transferring sensory inputs into memory. Here we might encounter a similar mechanism in which changes in population representations could lead to categorical responses protected devoid of sensory information.

Most previous studies have shown that the delay activity in sensory cortices is not causally involved in the behavioral response of trained animals (causal inactivations in V1^23^ and A1^25^). Note that differences of A1 firing rates between Go and No-Go sounds were not shown in studies with a delay^25,26^. However, delay activity may reflect postlearning residual top-down feedback, or alternatively may be the signature of an eligibility trace, i.e. a persistent activity necessary for bridging the gap between the sound and the response window^27^. Both scenarios are consistent with correlated categorical responses in the stimulus and delay periods. Based on our results, we interpret this delay activity in A1 during task engagement to be a category-based decision signal, as it is both conditioned by the objective category of the stimulus and exhibits dynamics that scale with the animal response times.

**Table-1:** Number of Units recorded.

| **Ferret** | **Recorded Units** | **Units used for**<br>**population analysis** | **Sessions** |
| --- | --- | --- | --- |
| Ferret P | 575 | 395 | 35 |
| Ferret T | 241 | 203 | 39 |

## Acknowledgments

We thank Joao Barbosa and Célian Bimbard for deep reading and fruitful comments. This work was supported by ANR-17-EURE-0017 and ANR-10-IDEX-0001-02, ERC 787836-NEUME to SAS, and ANR-JCJC-DynaMiC to YB.

## Methods

### Animals

Adult female ferrets (Mustela putorius furo) obtained from Marshall BioResource were used for this study. The animals were 1-3 years of age, weighing 500-800 g and were housed in pairs or trios with a normal day-night light cycle and free access to water during weekends. Ferrets were on a water-controlled protocol in which their water intake is restricted during the week days. Water was delivered during behavioral sessions as a reward. To maintain a stable weight, we provided ad libitum water for 1-2 hours post-behavior. The animals’ weights were daily monitored and maintained at 80% of pre-experiment weight.

### Behavioral task & training

Two adult female ferrets (*Mustela putorius furo*) were trained on a Go/No-Go delayed categorization task and one additional ferret is used for naive recording. Stimuli were click trains grouped in two behavioral categories: No-Go (4, 8, 12 Hz) and Go (16, 20, 24 Hz) sounds. Clicks were monopolar, rectangular pulses of 1 msec duration with amplitude set at 70 dB sound pressure level. A trial started with a pre-stimulus silence of 0.5 sec followed by a click train (Go or No-Go) of 1.1 sec. Following an appetitive Go/No-Go task paradigm, the animals were trained to wait for a response window that started after a delay of 1 sec following stimulus offset. A hit (lick in Go sound trials) on the response window for Go stimuli was rewarded with 0.2 ml of water. A LED attached to the water spout emphasized the delay period in which the animal had to restrain from licking. Early trials (lick during stimulus and delay period) and false alarms (lick on the response window of a No-Go sound) were punished with a time-out of 10 sec. In each session, Go and No-Go stimuli were presented in a pseudo-random manner. In the absence of delay, ferrets learned to associate the categories in one week and we then slowly increased the delay between stimulus offset and response window. It took several weeks for the ferrets to be trained on the full task structure. Initially we trained with extreme categorical stimuli (4 and 24 Hz) that were easy to learn and after reaching a consistent performance (d’>1), we progressively introduced other stimuli.

### Surgery

To obtain stable neurophysiological recordings we implanted the ferrets with a stainless steel headpost. Day before the surgery, ferrets were injected with antibiotics (Baytril, 12.5 mg/kg subcutaneous) to minimize infections arising from the surgery. On the day of surgery ferrets were deprived of water and food 90 mn prior to the surgery. After sedation with medetomidine (0.08 mg/kg), anesthesia was induced with ketamine (5 mg/kg, intramuscular). Animals were kept under deep anesthesia (1-2% isoflurane) throughout the surgery and vitals (ECG, pulse, oxygenation and rectal temperature) were continuously monitored. We also medicated the animals with atropine sulphate (0.2 mg/kg) to stabilize salvation and control arrhythmia arising from anesthesia. Using a complete sterile procedure, the animal skull was surgically exposed by an incision to the skin along the media crest down to the neck. The temporal muscles were carefully removed from the medial crest to the beginning of the zygomatic arch and the lateral wing at the lateral end of the nuchal crest. Using a stereotaxic apparatus, the headpost was mounted on the skull using methyl methacrylate based dental adhesive resin cement. Stainless steel screws were anchored along the areas surrounding the auditory cortex leaving a cavity for easy access to the auditory cortex. Finally the surrounding areas are filled with poly-methyl methacrylate (PMMA) based bone cement to stabilize the implant. Antisedan, antidote to medetomidine, (0.4 mg/kg, SC), antibiotics (Baytril, 12.5 mg/kg, SC) and analgesic (meloxicam, 0.05 mg/kg, oral) were administered to the animal following the surgery.

We allowed a two-week post-operative care for the animals to recover from the surgery. Antibiotics were continued for 7 days and anti-inflammatory and analgesics were administered for 4 days. Animals were habituated to a head restrained custom made horizontal plastic tubes few days prior to training session. Animals were passed the ethical approval (authorization:01236.02) from the French Ministry of Agriculture and all experiments adhere to European directive guidelines (2010/63/EU).

### Neurophysiological Recordings

We chronically implanted 32-channel metal electrodes arrays (MEA; Pt-Ir, MicroProbes, 8 x 4, electrode impedance: 2.5 MΩ with 0.4 μm distance between the electrodes and 0.7 μm long) over the auditory cortex. We custom-designed the chronic implant with MEA inserted in a drive-shuttle system having a flexible control of the array vertical movement. The base of the drive is sealed with a stretchable silicon membrane sheet to stop flowing any residues into the drive. Before the implantation, the electrodes are moved down such that the apex popped out of the silicon membrane. Under surgical anesthesia (Isoflurane 1%), we made 4 mm x 4 mm craniotomies into the cement implant at locations defined during the surgery. These craniotomies allowed us to visually identify primary auditory cortex regions. Using a microscope, we carefully removed the transparent dura. This was to ensure that the array penetrated the brain without any strain from the dura. The drive-shuttle system was placed on the brain surface using a stereotaxic apparatus and the entire system was fixed to the skull using bone cement. To minimize the displacements due to the external movements, the chronic implant was enclosed in a custom-made tube cemented to the implant. Immediately after the surgery we slowly lowered the electrodes and observed physiological activity, allowing us to verify the electrodes moved inside the brain.

Each session of recording consisted of passive and active sessions. Recordings were performed head-fixed in a soundproof chamber. In the passive sessions the water spout was removed. Continuous electrophysiological recordings were digitized (31250 Hz), amplified (15,000 x) and band-passed between 300Hz and 7000Hz using a digital acquisition system (Blackrock Cereplex). Band-passed signals were monitored online and units (including multi-& single-) were identified by spikes crossing a threshold of 3 SD of baseline noise. The data acquisition was done using an open-source suite MANTA v. 1.0^28^. We used a custom made open-source software Behavioral Auditory PHYsiology (BAPHY) written in MATLAB for sound delivery, recording, behavioral monitor and online analysis.

To identify units, we presented band-pass noise (0.2 s duration, 1 octave bandwidth) and pure tone stimuli to the animal using earphones (Sennheiser IE 800). The primary auditory cortical responses were identified by analyzing tuning properties to 100 ms tone pips of random frequencies spanning 4 octaves and temporally orthogonal ripple combinations (STRF)^29^. A1 responses show sharp tuning to random tones and single peak, short latency STRFs^5,10,20^. Finally assessing the STRFs, we continued with the experimental protocols to record the neuronal responses.

### Unit identification & Spike Sorting

We performed offline sorting on thresholded signals using PCA-based customized spike-sorting routines written in MATLAB. Single and multi-unit responses were identified by spiking shape and manually adjusting the PCA clusters^10^. A total of 575 (Ferret P) and 241 (Ferret T) multi-units were identified and used for further analysis. Spike sorting was done on concatenated passive and active sessions. We obtained 11.3±4.9 neurons per session (±std; n = 35 sessions) for ferret P and 5.2±3.2 neurons per session (±std; n = 39 sessions) for ferret T.

### Data Analysis

Offline data analysis was performed using custom written scripts in MATLAB (R2016a). All units were pre-processed to identify stable units that were kept under recording for both the active and passive sessions. We used a firing rate-based threshold to find stable units with non-zero firing rate for > 80% of the trials and the difference between time-averaged maximum and minimum firing rates is less than 10 fold across trials. We only analyzed units with > 2 spikes/sec firing rate. This procedure yielded 395 units in ferret P and 203 units in ferret T. Spike counts were constructed for 100 ms nonoverlapping time bins and thus used for further analysis. All population analysis were done at the single-session level, and therefore individual sessions were used a samples in statistical tests. This also allowed us to use all trials in each session, despite the difference in the number of correct trials across sessions due to variable behavior.

*1.a Delay Sustained neurons* Neurons with sustained activity during the delay were identified using z-scored spike counts from the baseline period (500 ms before stimulus onset). We then applied a threshold of 1.5 to the average spike count during the delay period.
*1.bModulation Index* For each unit, the modulation index of activity for quantifying modulation of delay activity was computed using:

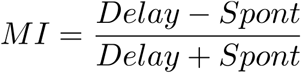

where *Delay* is the firing rate during the delay and *Spont* is the spontaneous activity.
*1.c Population decoding* In each recording session, we constructed binary linear classifiers to decode stimulus categories (Go vs No-Go) at each time step with 100 ms binning (n= 200 cross-validations). Classifier performance was statistically evaluated through permutation tests, comparing the actual performance with a chance-level distribution obtained with 200 label permutations (the lower bound for the p-value being 1/200 = 0.005). We opted for 4-fold cross-validation. We balanced trial number across categories. Unless mentioned otherwise, all analysis were done using only correct trials (correct rejection No-Go trials and hit Go trials). Temporal evolution of the decoder was computed as the correlation between decoding weight vectors at one time bin against others.
*1.d Linear Regression* A linear decoder trained on classifying categories inherently mixed sensory and category information to decode categories. To disentangle the contributions from sensory features and categories in the population code, we opted for linear regression models to capture the unique contribution of each feature. The linear model was constructed by combining sensory click rates and binary values (−1 and +1) for categories into a design matrix, to capture the contribution of different task variables. We used ridge regression to avoid overfitting. The ridge parameter is calculated algorithmically using cross-validation ^30^. The regression model was as follows

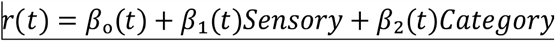 Chance level was obtained through permutation tests (100 label permutations; the lower bound for the p-value being 1/100 = 0.001).
*1.e CPD* To capture the contribution of each feature into unit responses, we computed a coefficient of partial determination (CPD) as the fraction of variance lost by shuffling one of the feature with respect to the full original model. Doing so, CPD captured the unique contribution of that feature (Musal). We thus fitted reduced linear models with one of the variables being shuffled in the design matrix in order to destroy the contribution arising from that particular task variable. We used 5-fold cross-validation to compute the mean square error (MSE) for the full and reduced models. CPD was defined as:

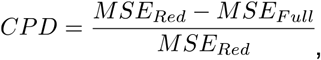

where *MSE_Red_* is obtained by cross-validating the following reduced models with 200 shuffles of the targeted variable:

- for estimating the unique contribution of sensory regressor:

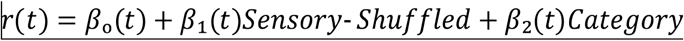
- for estimating the unique contribution of category regressor:

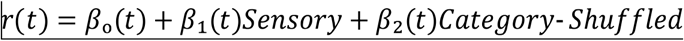 Time windows for computing average CPD in each period were 0-1.1 s for the stimulus period, 1.1-2.1 s for the delay period, and 2.1-3.1 s for the response window period.
*1.f Categorical and sensory population responses* We assessed the time-evolution of population activity along sensory- and category-related neural axes defined by the linear regression. This was done by projecting baseline-corrected population activity specific to the feature of interest onto category or sensory regressor weights. Therefore any deviation from zero represents the deviation of population activity along the regression axis away from the projection of baseline activity. Projections of trial-averaged population activity onto the category or sensory regressors were performed by subtracting the contribution of the other feature, i.e 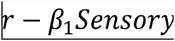 was projected onto the category and 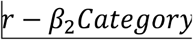 was projected onto sensory regressors trained at appropriate time epochs. We defined categorical (resp. sensory) projections as projections on the category (resp. sensory) axis. We could then project population activity corresponding to individual click rates (Fig 3a-d) or averaging across click trains of the same category (Fig 3e and Fig 4). Time windows for computing average projection axis in each period were 0.2-0.9 s for the stimulus period, and 1.3-2.1 s for the delay period.
*1.g Categorization Index* We quantified the amount of category information present in the projections by using categorization index (CI). CI is defined as a modulation index for the distance between and within categorical responses:

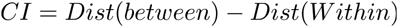
*CI = Dist(between) — Dist(Withiri*) CI was computed using all successive pairs within (4-8 Hz, 8-12 Hz, 16-20 Hz and 20-24 Hz) a category and one between category pairs (12-16 Hz).
*1.h Statistics* We performed both population decoding and linear regression analysis session wise including simultaneously recorded neurons. All statistics across passive and active states were done with t-tests while computing significance for each state were done through permutation tests. Unless specified otherwise, error bars showed ± 1 sem over sessions.

## Supplementary Figures

**Fig S1:**
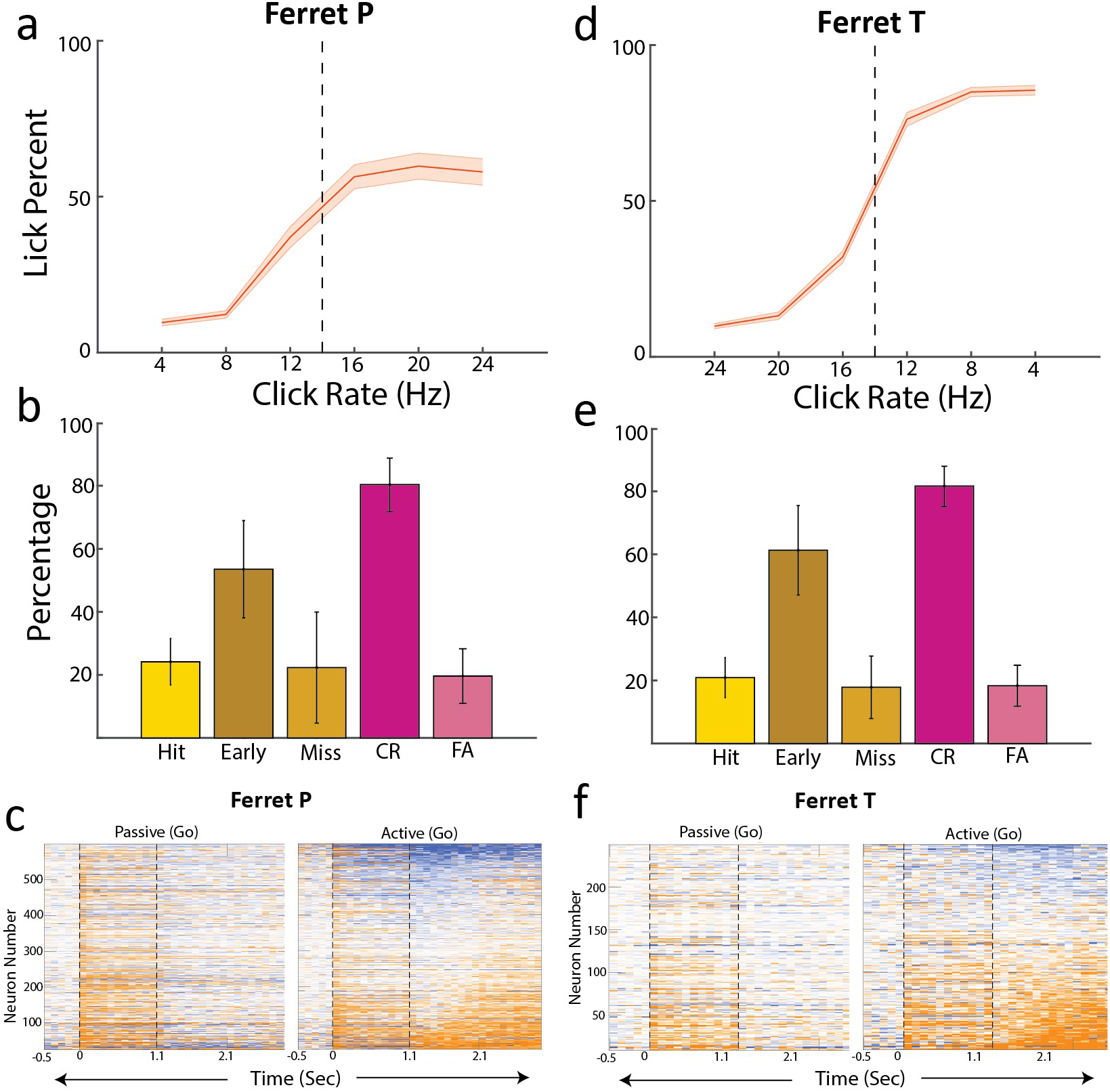
Behavioral performance and sustained responses in all animals. Behavioral performance and sustained responses for all animals. **a.** Psychometric curves. **b.** Average percentage of trials in recording sessions. **c.** Sustained activity for Ferret P. **d,e,f.** Same as **a,b** and **c** for Ferret T.

**Fig S2:**
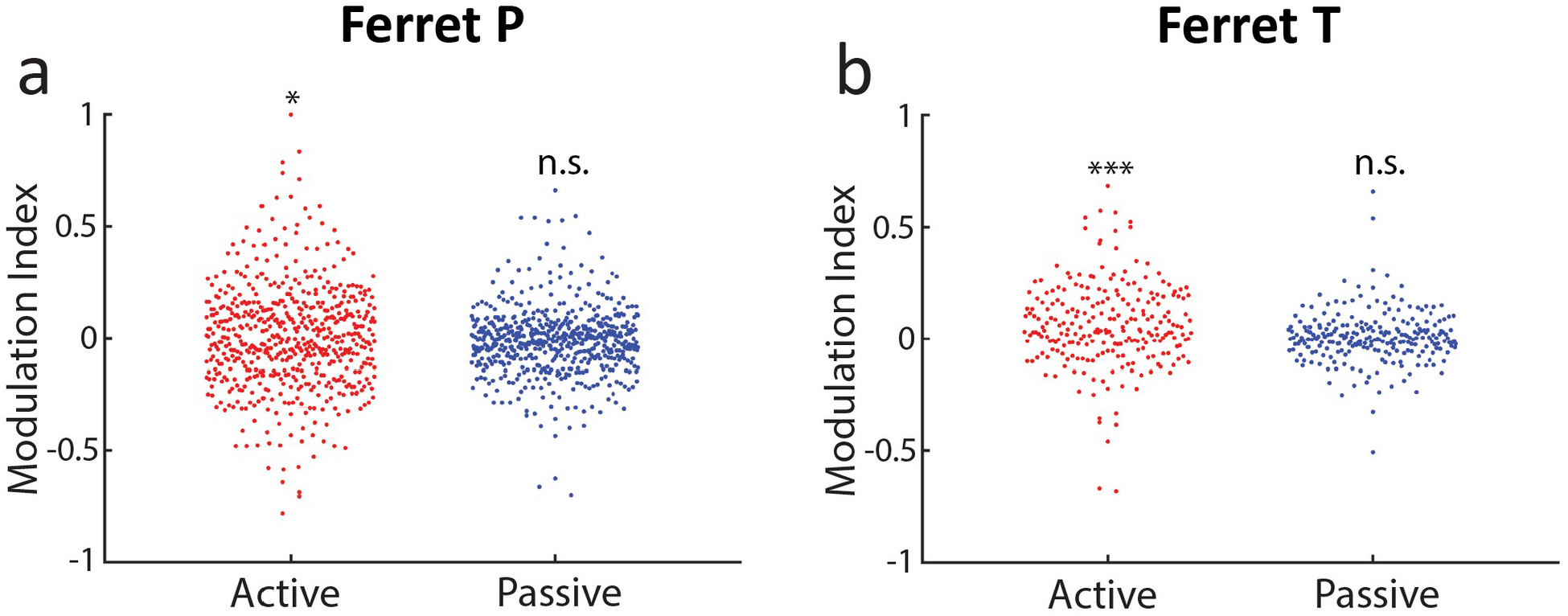
Percentage of neurons in all animals. Modulation Index for delay activity in individual ferrets (*** p<0.001, * p<0.1).

**Fig S3:**
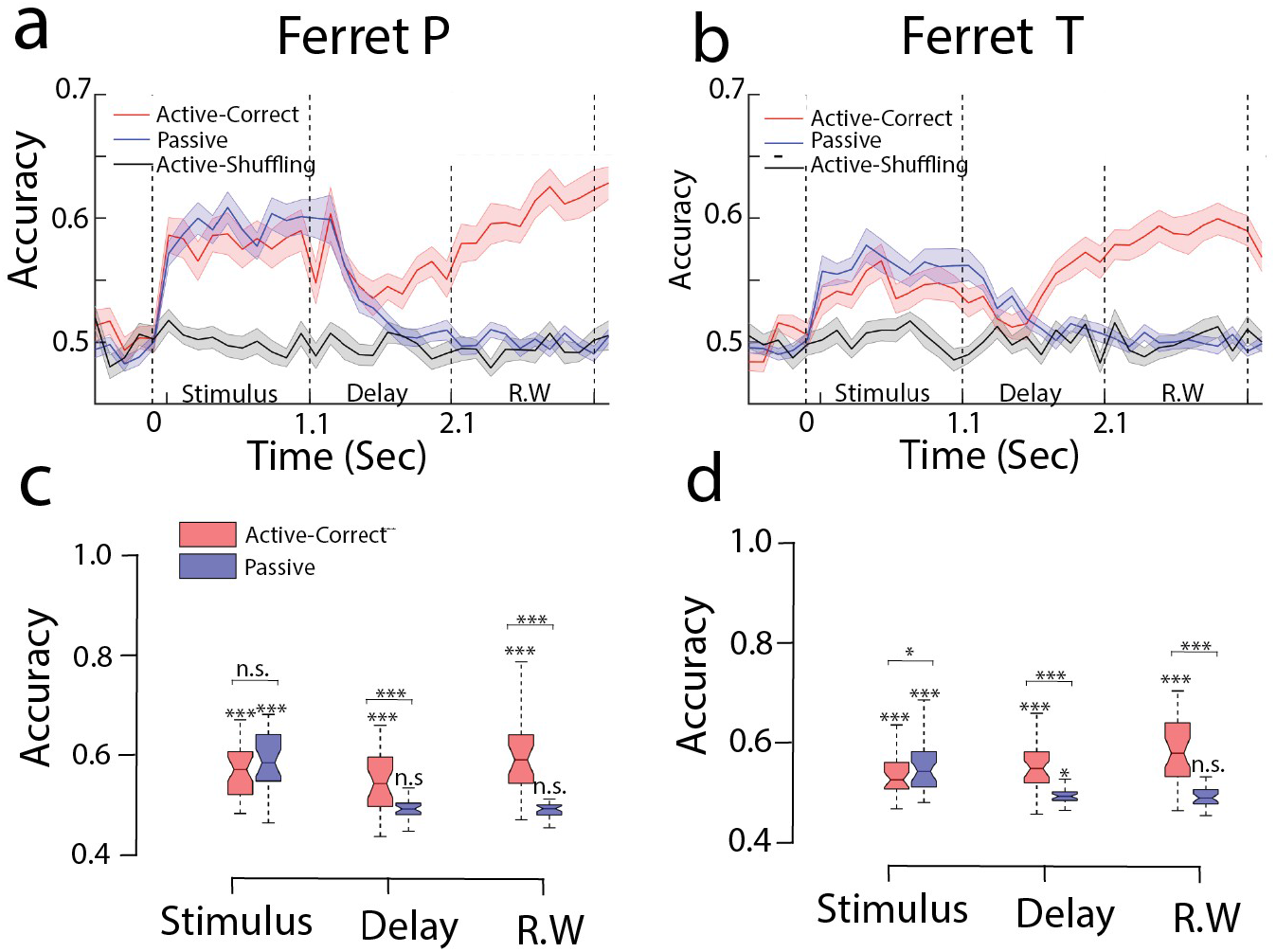
Decoding accuracy split by animals. Decoding accuracies for passive, active-correct behavior for individual ferrets (Ferret P: n=35 sessions, Ferret T: n=39 sessions).

**Fig S4:**
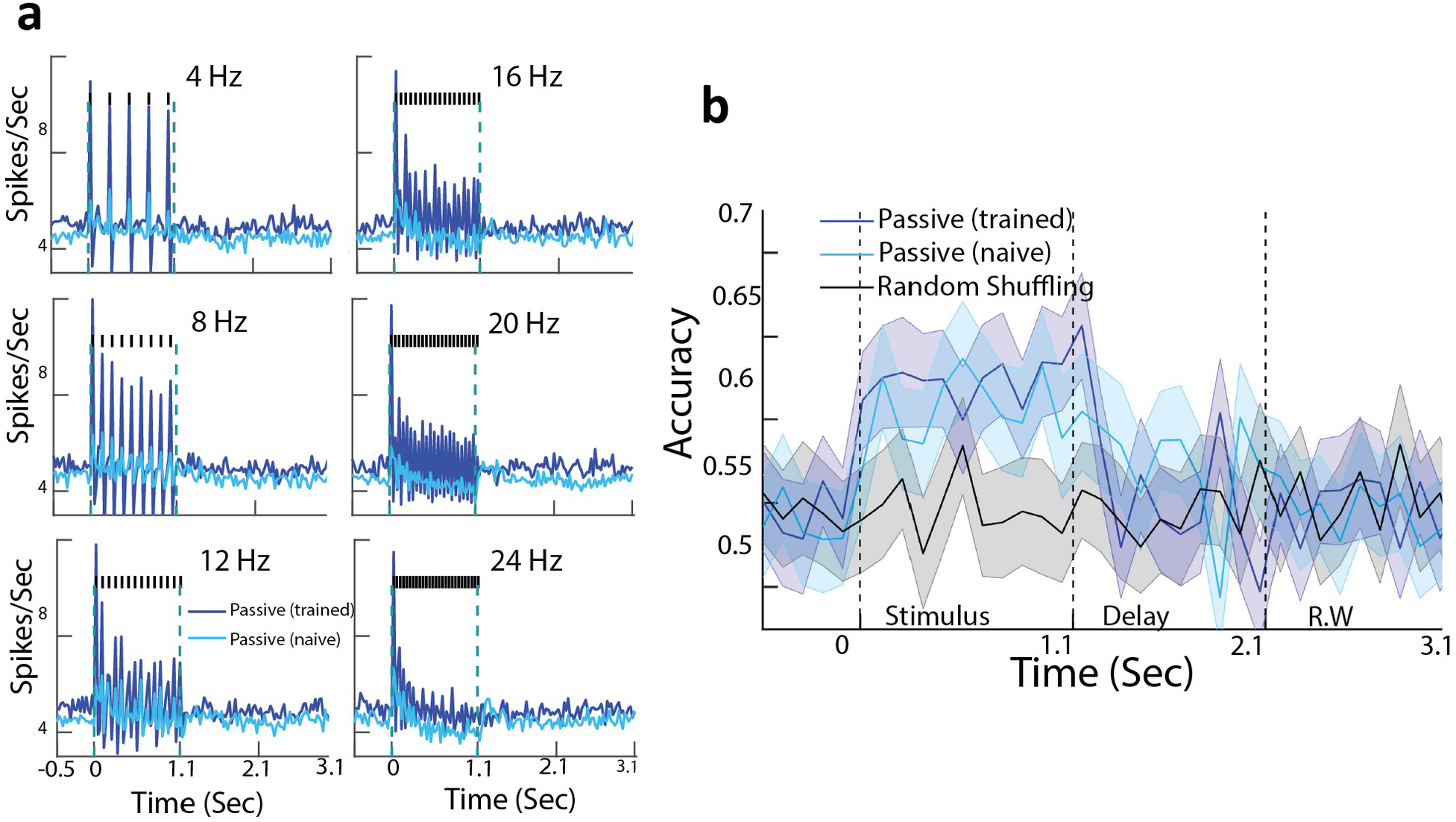
A1 Population encoding in naive animal. Decoding accuracy in naive animals. **a.** Population averaged PSTHs for passive listening and (deep blue) and in naive animals (light blue). **b.** Decoding performance for naive animals. Gray curve indicated the performance by shuffling labels in task engagement. Error bars are 1STD over 200 crossvalidations. Neuron number is balanced while computing the decoders for trained and naive animal (n=71 units).

**Fig S5:**
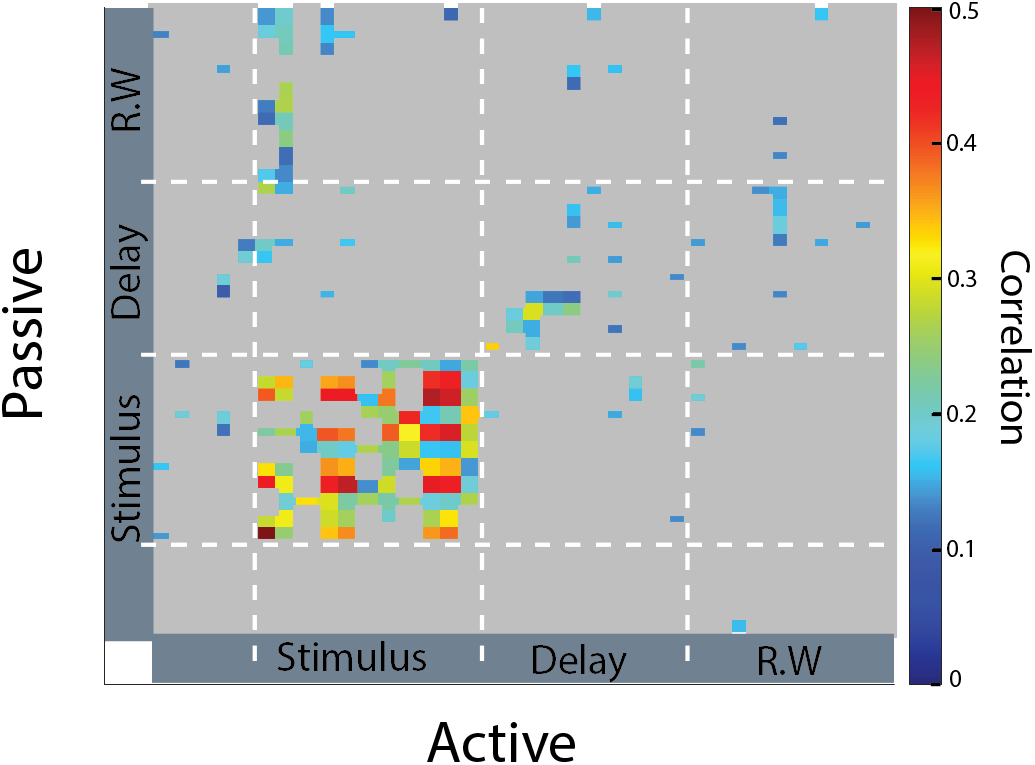
Correlation of passive and task-engaged decoders. Correlation of passive and task-engaged decoders for Ferret P.

**Fig S6:**
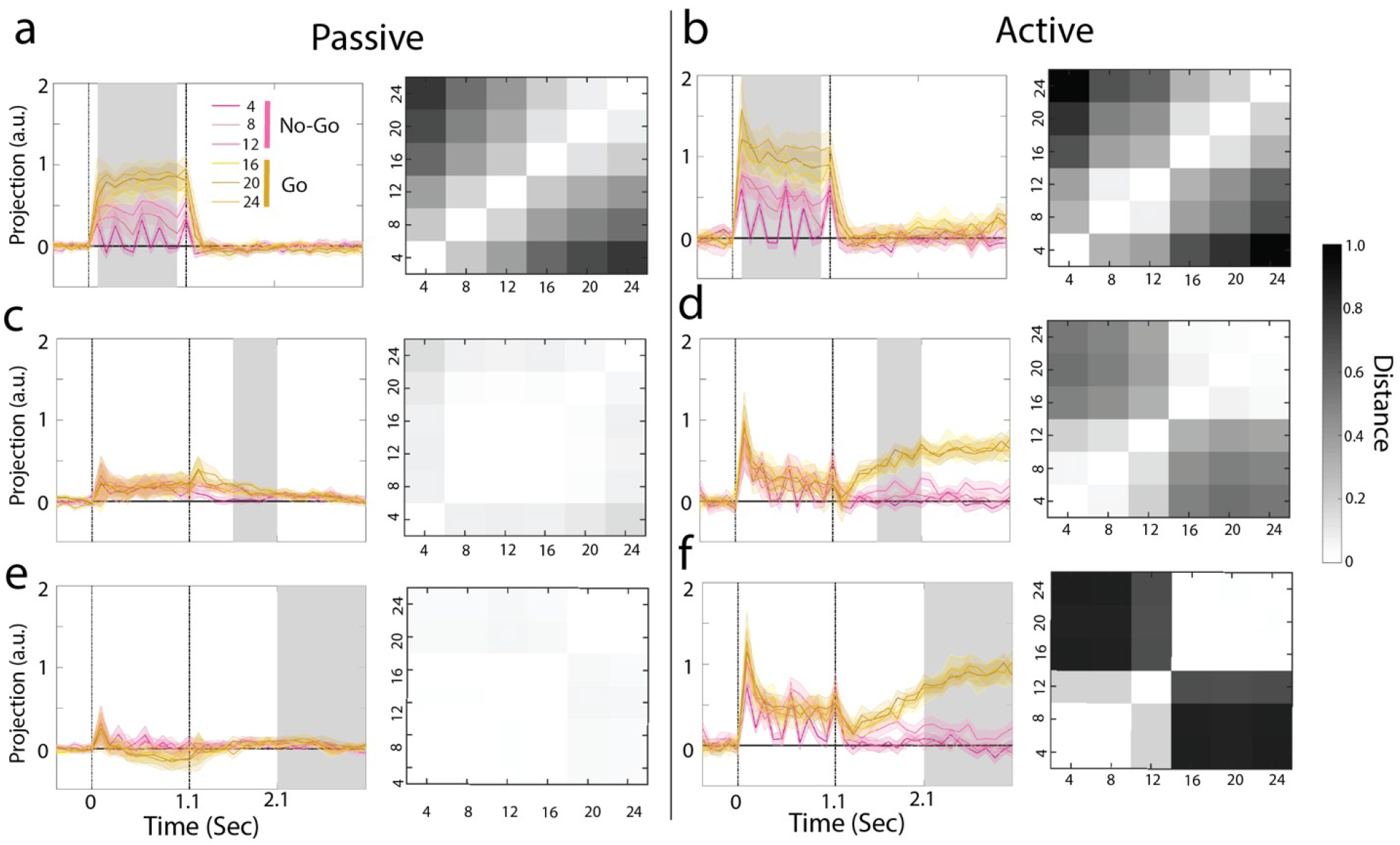
Projection of each click rates onto linear decoders. **a,c,e.** Projection of trial averaged activity of each of the category stimuli onto the passive decoders trained at different time epochs (i.e., stimulus, delay and response window). The gray shaded region indicates the time at which the decoder was trained. Right panels show the distance matrix between successive stimulus pairs. **b,d,f.** Same as **a,c,e** for active behavior.

**Fig S7:**
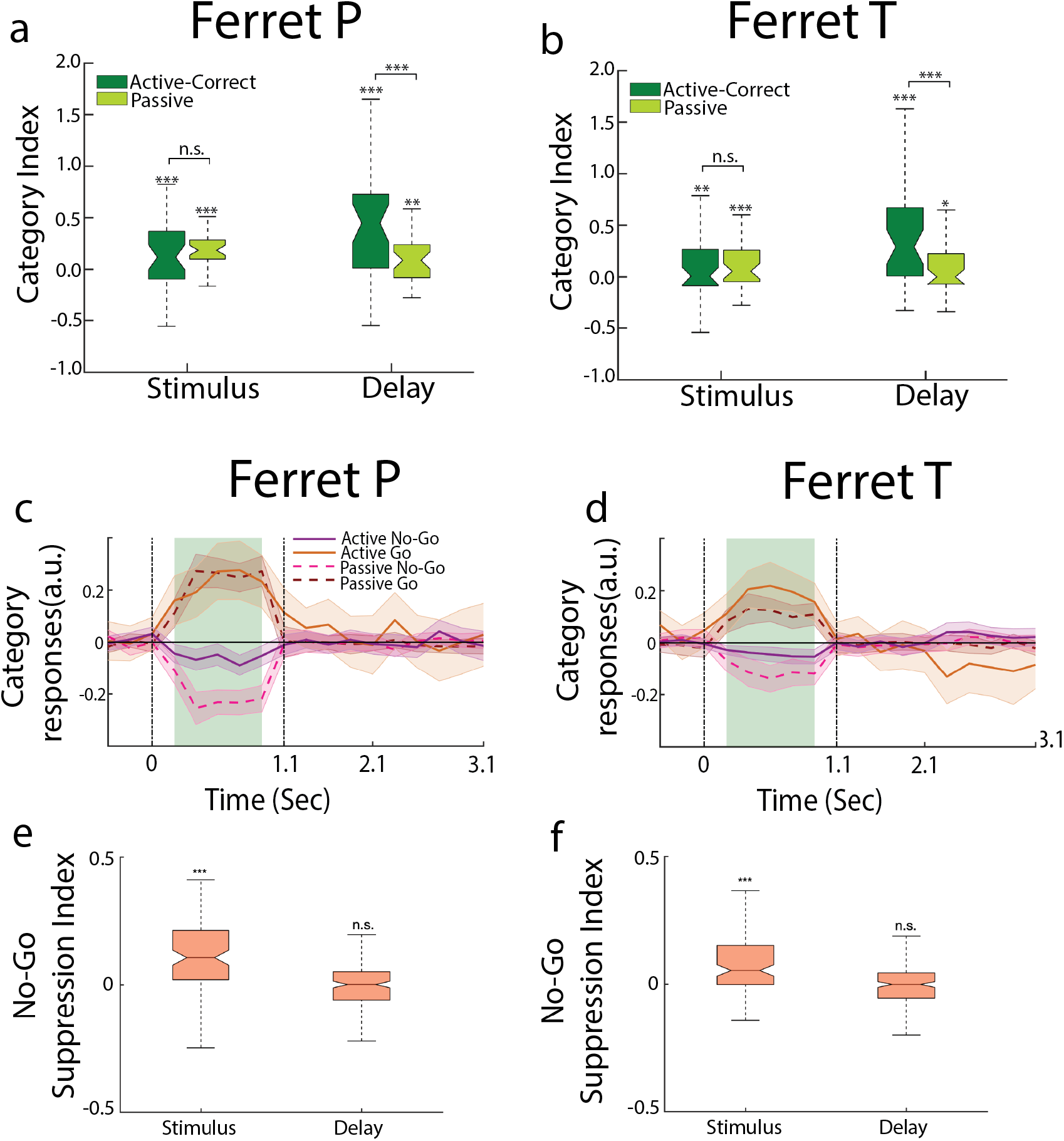
No-Go Suppression index from the projections onto category regressor during sound period split by animals. No-Go Suppression in all animals. **a,b.** Categorization Index for ferret P and ferret T. **c,d.** Projection of trial average activities of passive and active states onto the stimulus category regressors for ferret P (c) and ferret T (d). **e,f.** No-Go suppression index.

**Fig S8:**
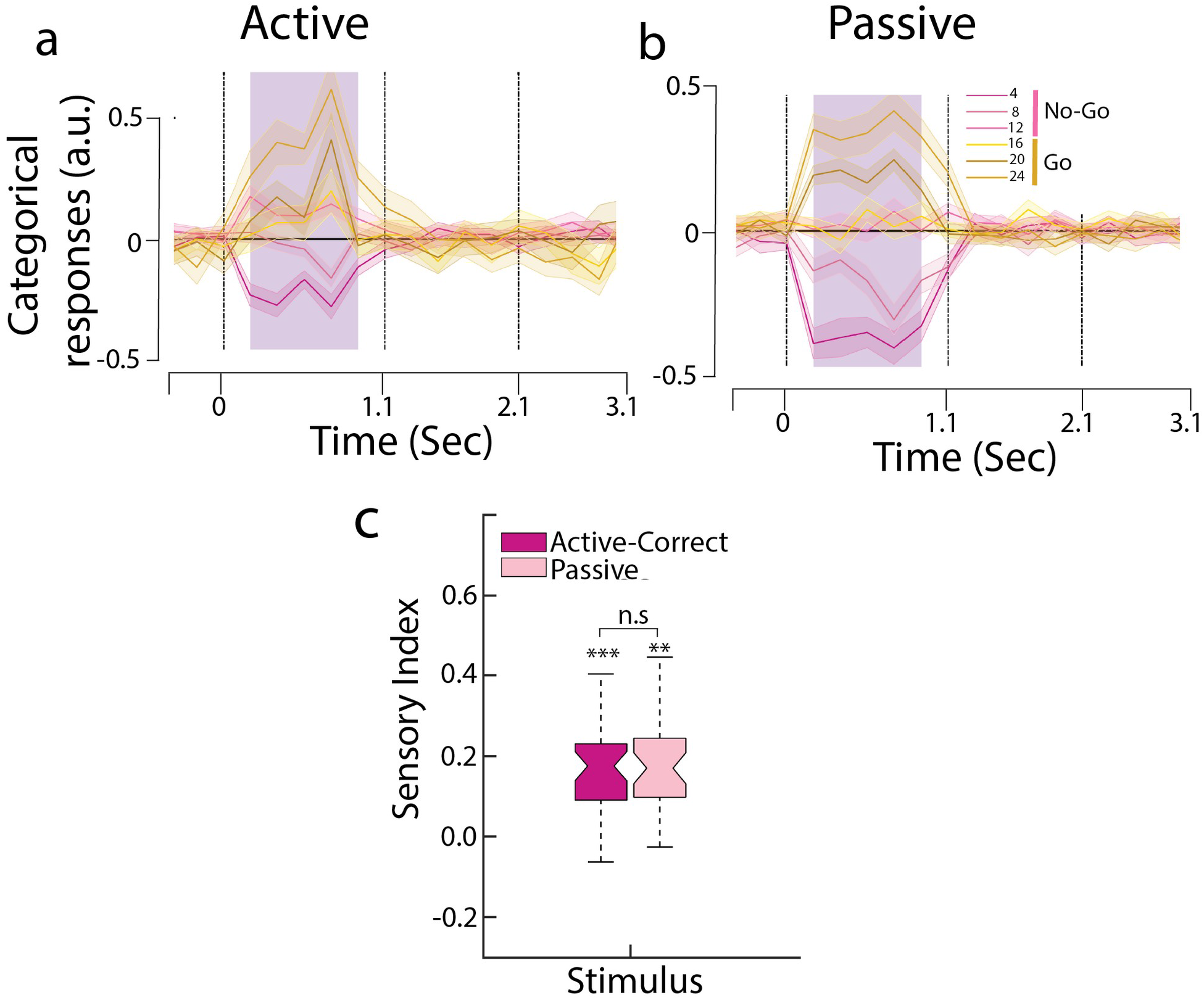
Projection of trial averaged activity onto sensory regressors. **a,b.** Projection of trial averaged activity onto sensory regressor trained at stimulus time epoch. **a** for active and **b** for passive state. **c.** Sensory index as quantified by projection distance between successive click rates on to the regressor (*** p<0.001 using permutation test for individual boxes).

**Fig S9:**
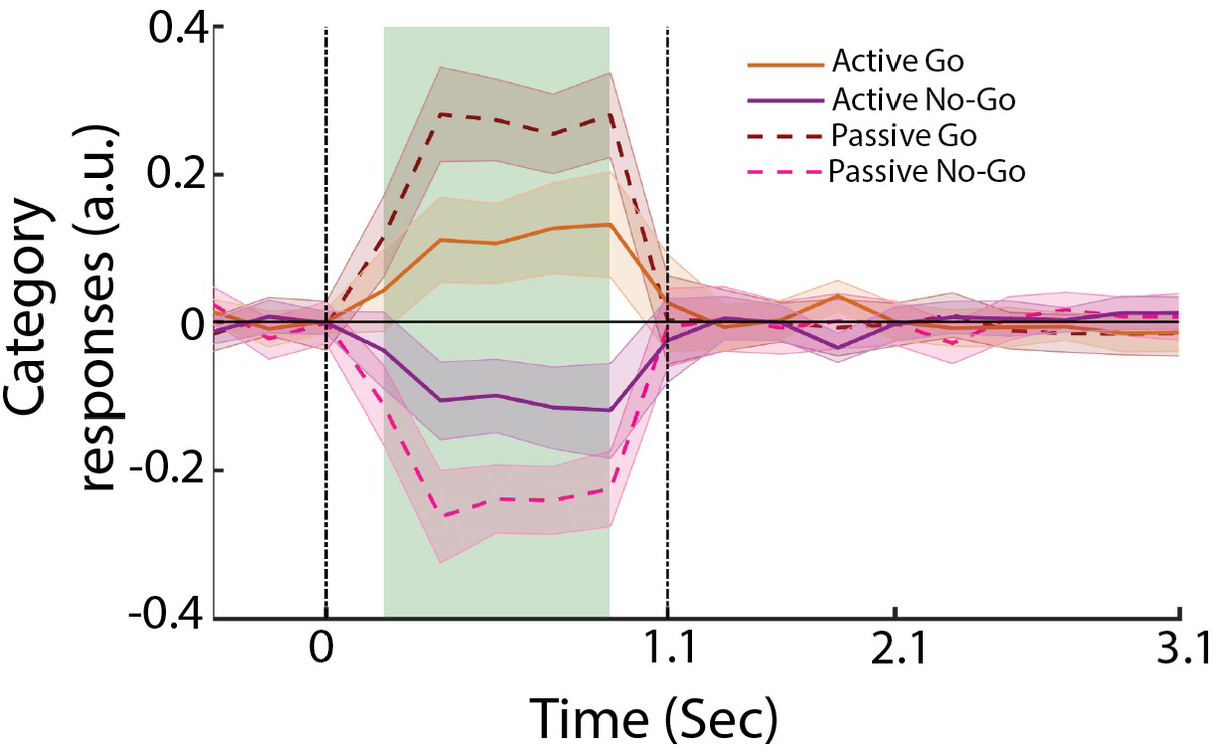
Projection onto category regressor extracted from the passive condition. Projection onto category regressor extracted from the task-engaged condition for passive and active states. Note that Go and No-Go sounds are symmetrically laid-out around baseline projections.

**Fig S10:**
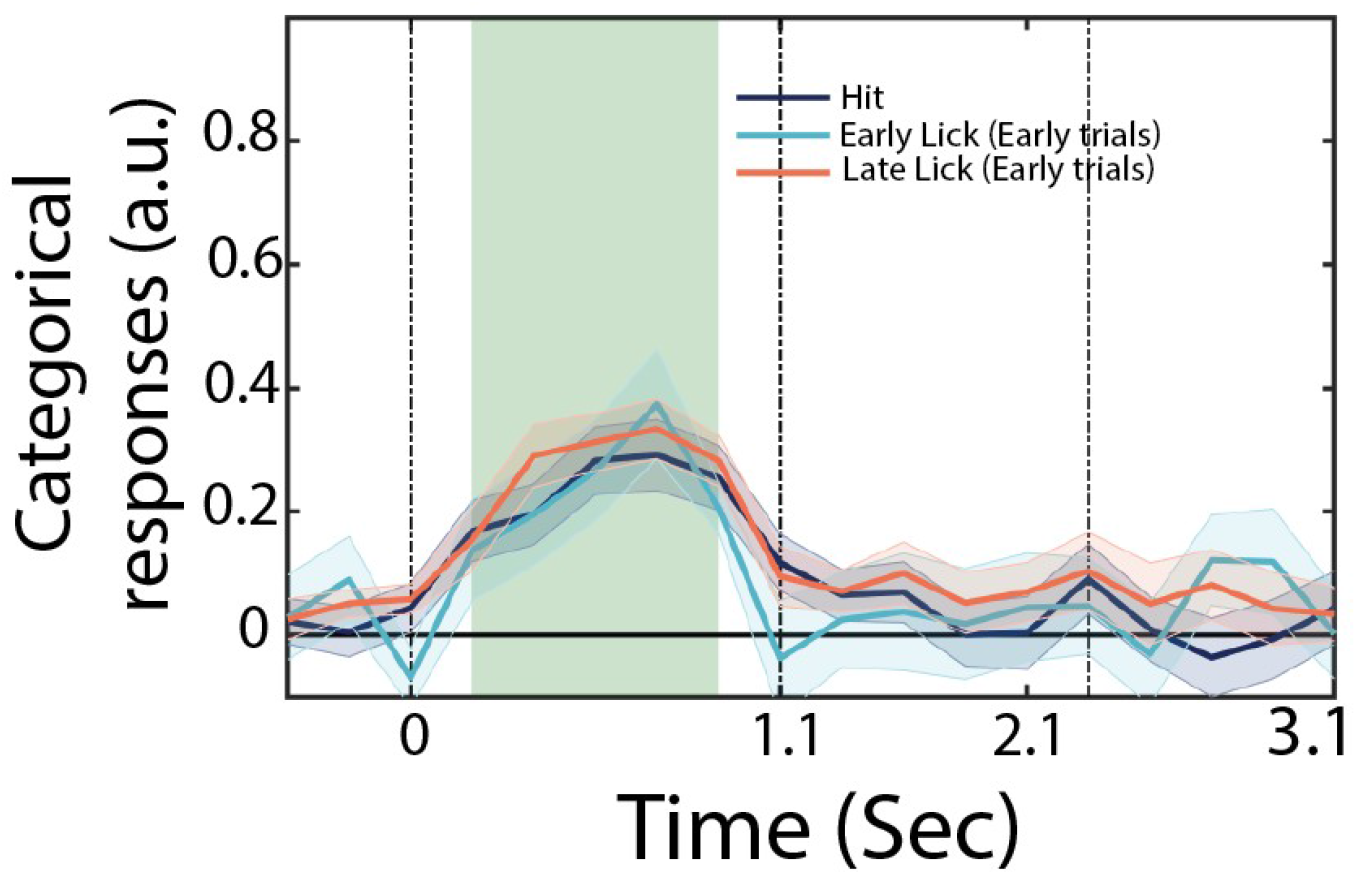
Like Fig 4c with categorical responses of the stimulus period. Projection of trial averaged activity of Hit, Early-Early and Late-Early trials onto the stimulus category regressor.

**Fig S11:**
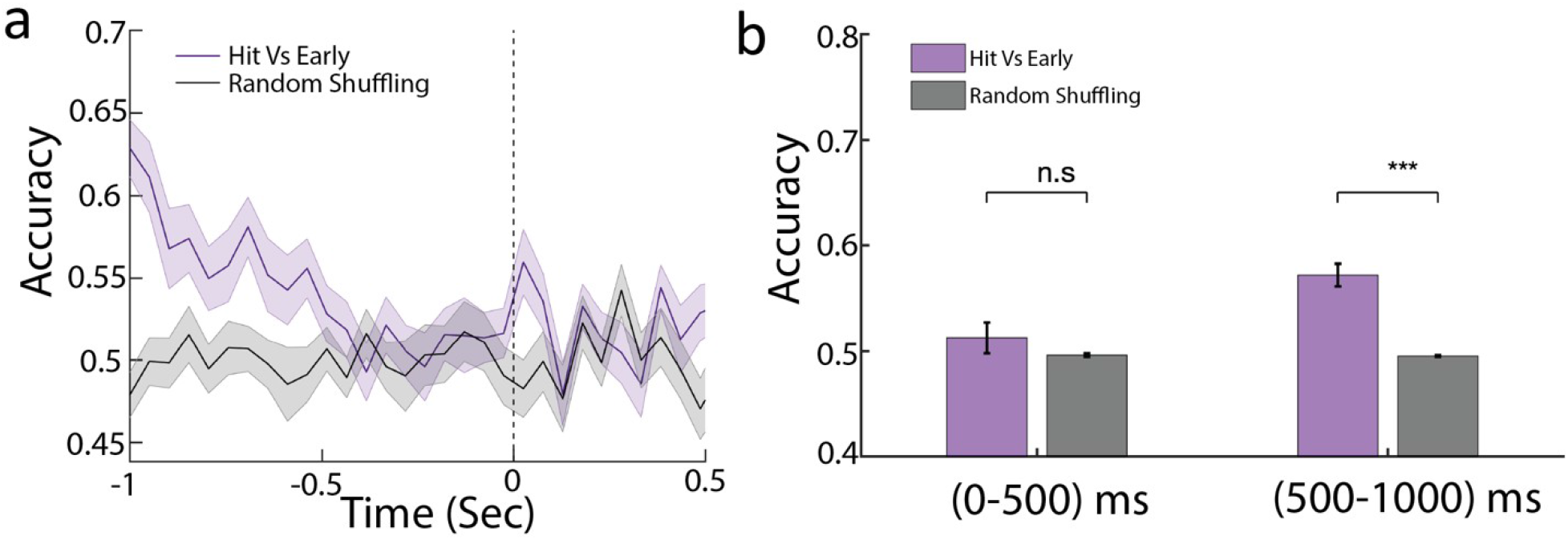
Delay activity in hit and early trials is identical. **a.** Lick-aligned population decoding of hit vs CR trials. The accuracy is about chance level 500 ms before the lick. **b.** Comparison of accuracy within 0-500 ms and 500-1000 ms before lick with that of random shuffling, significance is computed using a permutation test (*** p<0.001).

**Fig S12:**
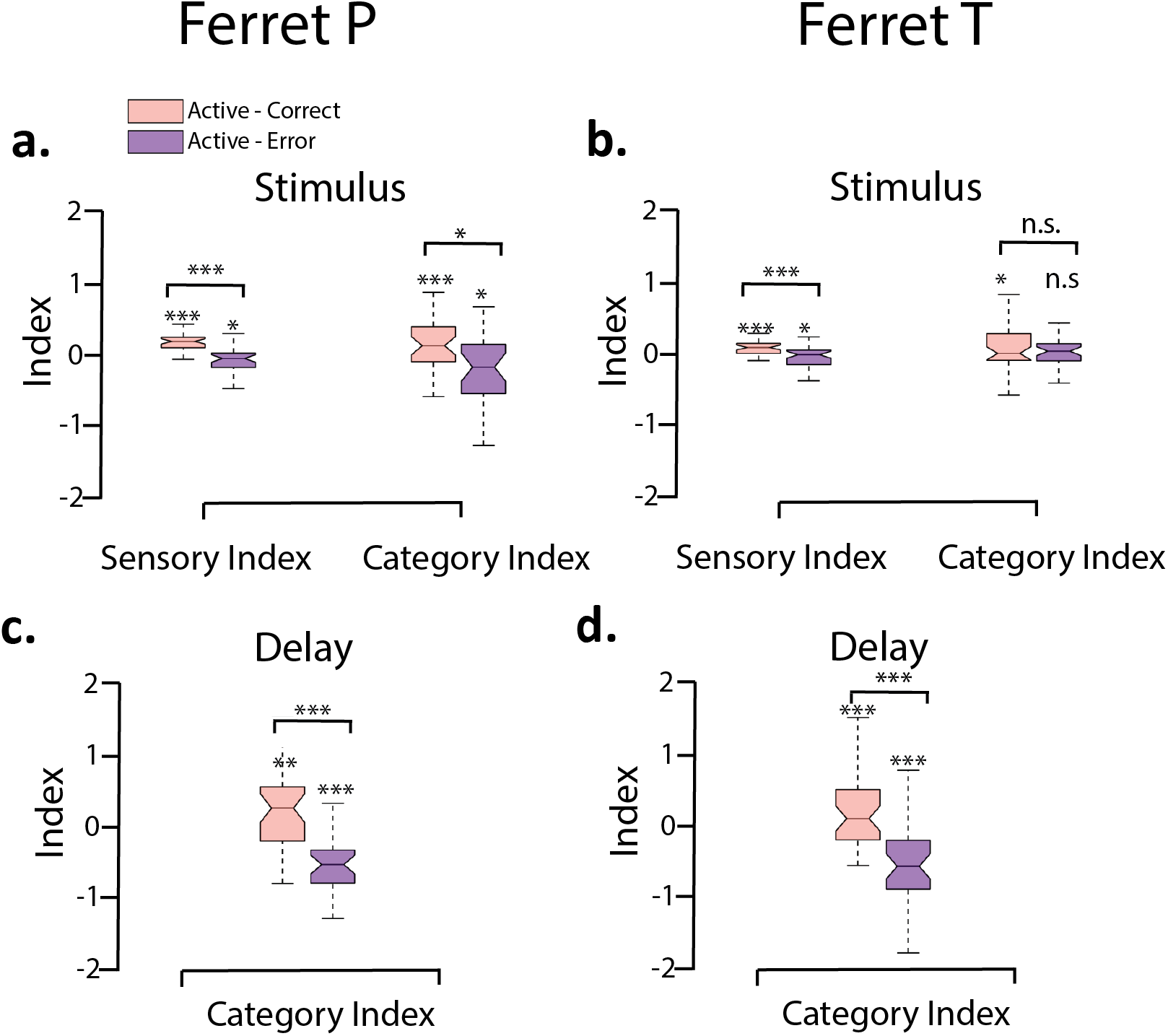
Sensory and Category Index split by animals. **a,c.** Sensory and Category Index during active correct and error trials for ferret P. **b,d.** same as **a,c** for ferret T (*** p<0.001, ** p<0.05, * p<0.1).

